# Reactive/proactive aggression specific cortical and subcortical alterations in children and adolescents with disruptive behavior

**DOI:** 10.1101/490086

**Authors:** Jilly Naaijen, Leandra M Mulder, Shahrzad Ilbegi, Sanne de Bruijn, Renee Kleine-Deters, Andrea Dietrich, Pieter J Hoekstra, Jan-Bernard C Marsman, Pascal M Aggensteiner, Nathalie E Holz, Boris Boettinger, Sarah Baumeister, Tobias Banaschewski, Melanie C Saam, Ulrike M E Schulze, Paramala J Santosh, Ilyas Sagar-Ouriaghli, Mathilde Mastroianni, Josefina Castro Fornieles, Nuria Bargallo, Mireia Rosa, Celso Arango, Maria J Penzol, Julia E Werhahn, Susanne Walitza, Daniel Brandeis, Jeffrey C Glennon, Barbara Franke, Marcel P Zwiers, Jan K Buitelaar

## Abstract

**Objective:** Maladaptive aggression, as present in conduct disorder (CD) and, to a lesser extent, oppositional defiant disorder (ODD), has been associated with structural alterations in various brain regions, such as ventromedial prefrontal cortex (vmPFC), anterior cingulate cortex (ACC), amygdala, insula and ventral striatum. Although aggression can be subdivided into reactive and proactive subtypes, no neuroimaging studies have yet investigated if any structural brain alterations are associated with either of the subtypes specifically. Here we investigated this association in predefined regions of interest.

**Method:** T1-weighted magnetic resonance images were acquired from 158 children and adolescents with aggressive behavior (ODD/CD) and 96 controls in a multi-centre study. Aggression subtypes were assessed by questionnaires. Cortical volume and subcortical volumes and shape were determined using Freesurfer and the FMRIB integrated registration and segmentation tool. Associations between volumes and continuous measures of aggression were established using multilevel linear mixed effects models.

**Results:** In cases only proactive aggression was negatively associated with amygdala volume (*b*=−11.82, *p*=0.05), while reactive aggression was negatively associated with insula volume (*b*=−46.41, *p*=0.01). Classical group comparison showed that children and adolescents with aggressive behavior had smaller volumes than controls in (bilateral) ventral striatum (*p*=0.003), ACC (*p*=0.01), and vmPFC (*p*=0.003) with modest effect sizes.

**Conclusions:** Aggression was associated with reduced volume in brain regions involved in decision making. Negative associations were found between reactive aggression and volumes in regions involved in threat responsivity and between proactive aggression and regions linked to empathy. This provides evidence for aggression subtype-specific alterations in brain structure.

## Introduction

Aggression, overt behavior with the intention of inflicting damage, is a behavioral trait with important roles throughout evolution in defense and predation. However, when expressed in humans in the wrong context, aggression may lead to social maladjustment and crime. Maladaptive aggression is commonly observed across childhood in disruptive behavioral disorders, in particular in conduct disorder (CD) and to a lesser degree in oppositional defiant disorder (ODD). CD is defined as a repetitive and persistent pattern of behavior, which violates the rights of others and major age-appropriate societal rules. ODD is characterized by a frequent and persistent pattern of irritable and angry mood, vindictiveness, and inappropriate, negativistic, defiant, and disobedient behavior toward authority figures.^1^ Both disorders are highly comorbid with attention-deficit/hyperactivity disorder (ADHD), which has been associated with aggressive behavior as well.^2^

Aggression is a heterogeneous phenomenon and has been subtyped in diverse ways. Such subtyping is considered an important step towards effective prevention and treatment strategies, as currently available treatment options for maladaptive aggressive behavior have limited efficacy.^3,4^ A promising subdivision, derived from animal studies, defines impulsive and instrumental subtypes of aggression, also referred to as reactive and proactive aggression, respectively.^5^ Reactive aggression is associated with high arousal, impulsivity, high affect and uncontrolled behavior. Animal studies have shown that this form of aggression is mediated by a circuit that is responsive to threat (and frustration) and involves the amygdala.^6^ Furthermore, this circuit may be regulated by frontal cortical regions, such as the ventromedial prefrontal cortex (vmPFC) and the anterior cingulate cortex (ACC).^7^ In contrast, proactive aggression refers to goal-directed, planned behavior associated with low arousal and is associated with higher levels of callous unemotional and/or psychopathic traits. This form of aggression often goes hand in hand with impaired stimulus-reinforcement learning (which involves the amygdala) combined with impaired prediction error signaling (which involves the striatum), leading to a poor understanding of the value of objects, cues and responses represented in the vmPFC^7^ as well as a lower empathy level. Although the subdivision of reactive versus proactive aggression is the most prevalent subdivision referred to in literature, no neuroimaging studies have yet investigated if any structural brain alterations are associated with either of these aggression subtypes specifically in a clinical sample.^8^ Prior research has focused on the presence or absence of callous unemotional (CU) traits and/or childhood versus adolescent onset of CD in subtyping aggression. The strongest evidence from such studies so far points to an involvement of the fronto-limbic-striatal circuitry in aggressive behavior.^7^

Several studies have focused on structural abnormalities related to aggression. Almost all of these performed voxel-based morphometry (VBM) analyses of grey matter.^9–12^ A recent meta-analysis of thirteen VBM studies included almost 400 participants (aged 9-21 years) with conduct problems and showed that individuals with conduct disorder/problems compared with controls had smaller grey matter volumes in the left amygdala, in the bilateral insula extending to the ventrolateral prefrontal cortex (PFC)/orbitofrontal cortex (OFC) and in the medial superior frontal gyrus extending to the anterior cingulate cortex (ACC) with small-medium effect sizes.^13^ Another meta-analysis including ODD/CD and ADHD studies (n=415, age 8-21 years) reported reduced volumes of the amygdala, insula and frontal regions in ODD/CD as well, and with greater reductions in the presence of comorbid ADHD.^14^ However, several other studies have not been able to find any group differences in grey matter (GM) volume in these regions between participants with conduct problems and controls.^15,16^ Furthermore, opposite results have been shown as well. Cohn (2016) reported a positive association between CU traits and GM volume in the insula and a positive association of CD symptoms with amygdala GM volume.^9^ Structural brain alterations were found to be accompanied by more consistently reported functional alterations associated with deficient empathy, heightened threat sensitivity, and/or deficient decision making and response inhibition in the amygdala, vmPFC and striatum as well.^17^

In the present multi-center study we investigated the association between structural alterations and continuous measures of reactive and proactive aggression as well as CU traits and ADHD symptoms in the largest sample of children/adolescents with disruptive behavior disorders reported so far. We used pre-selected regions of interest based on the previous meta-analyses and thus investigated ACC, insula, lateral and medial parts of the orbitofrontal cortex (referred to as vmPFC), amygdala, and ventral striatum and expect their volume to be negatively associated with re- and/or proactive aggression. We chose not to perform a whole brain VBM analysis to limit the number of independent tests^18^ and because surface-based morphometry measures have been shown to be more robust across different MR scanners, as used in our multi-site design.^19^ We focused on analyses of the volumes of these regions and subsequently investigated the shape of the subcortical structures for subtler morphological changes and cortical thickness and surface area of the cortical areas.

## Methods and materials

### Participants

We included 277 participants (n=176 cases and n= 101 healthy controls) aged 8-18 years who were recruited across nine sites in Europe (see supplemental material for details). Exclusion criteria for all participants were contraindications for MRI, an IQ<80 and a primary DSM-5 diagnosis of psychosis, bipolar disorder, major depression and/or anxiety disorder. Participants that were included as “cases” were diagnosed with conduct disorder (CD) and/or oppositional defiant disorder (ODD) or scored above the clinical cut-off for aggressive behavior and/or rule-breaking behavior as measured with the Child Behavior Checklist completed by parents (CBCL).^20^ In the healthy comparison group, no DSM axis I disorder was allowed in the participants. Participants that were using medication were at a stable dose for at least two weeks. Ethical approval for the study was obtained for all sites separately by local ethics committees. After description of the study written informed consent was given by the participants and/or their parents.

### Phenotypic information

Diagnoses of ODD, CD and/or ADHD were confirmed by structured diagnostic interviews with both child and parents using the Kiddie Schedule for Affective Disorders and Schizophrenia (K-SADS).^21^ Participants were administered a screening-module, followed, if needed, by application of disorder-specific modules. Aggressive behavior was measured by the aggressive behavior and rule-breaking behavior sub-scales of the CBCL.^20^ The Reactive Proactive Aggression Questionnaire (RPQ)^22^ completed by participants themselves was used to subtype aggressive behavior. The presence of CU traits was assessed by the Inventory of Callous Unemotional Traits (ICU)^23^ completed by parents. A continuous measure for ADHD symptoms was derived from the K-SADS by summing the number of inattention and hyperactivity/impulsivity symptoms. IQ was estimated from four subtests (vocabulary, similarities, block design and picture completion/matrix reasoning) of the Wechsler Intelligence Scale for Children III or IV.^24^ Information about use of medication was collected via parental report on the measurement day.

### MR acquisition and processing

MRI data-sets were acquired on 3T scanners across nine different sites in Europe (see **Table 1** for the scan parameters). The T1-weighted images were processed with the FMRIB Software Library (FSL)^25^ for subcortical volume and shapes and with Freesurfer v5.3.0 (http://surfer.nmr.mgh.harvard.edu) for measures of cortical volumes and cortical thickness (CT) and surface area (SA).^26,27^ Subcortical segmentation was performed with the automated FMRIB integrated registration and segmentation tool (FIRST)^28^ which included affine registration to MNI space followed by a segmentation procedure that integrates both shape and intensity information for accurate segmentation of subcortical structures, including the bilateral ventral striatum and amygdala. Volumes of these respective regions were extracted for statistical analysis. Vertex analysis was performed with FIRST_utils to determine shape. A multivariate Gaussian model of the location and intensity variation of the vertex was used to generate surface meshes. Localized shape differences using the 3D-coordinates of the corresponding vertices with vertex-wise F-statistics were calculated after alignment to the average shape of the cohort and removal of global scaling (useRigidAlign and useScale). Cortical reconstruction was performed in FreeSurfer using the Desikan Killiany atlas (see **Figure 1**). CT was calculated for each vertex on the reconstructed cortical sheet and was defined as the closest distance between the grey/white matter boundary and the grey matter/CSF boundary.^29^ SA was measured at the geometric middle of the inner and outer cortical surfaces. CT, SA and volume of the ACC (rostral and caudal labels), vmPFC (medial orbitofrontal and lateral orbitofrontal labels) and insula were extracted from the parcellations and segmentations using the ‘mri_segstats’ function.

**Table 1.** Scan parameters for the T1-scan across the different sites

| Scanner | Site | TR*/TE/T1<br>(ms) | Flip<br>angle | Field of<br>view | Matrix<br>RL/AP/slices | Voxel size<br>(mm) | Acceleration<br>factor |
| --- | --- | --- | --- | --- | --- | --- | --- |
| **Siemens** | Nijmegen | 2300/2.98/900 | 9 | 256 | 212/256/176 | 1.0x1.0x1.2 | 2 |
|  | Mannheim | 2300/2.96/900 | 9 | 256 | 212/256/176 | 1.0x1.0x1.2 | 2 |
|  | Ulm | 2300/2.96/900 | 9 | 256 | 212/256/176 | 1.0x1.0x1.2 | 2 |
|  | Barcelona | 2300/2.98/900 | 9 | 256 | 212/256/176 | 1.0x1.0x1.2 | 2 |
|  | Madrid | 2300/2.98/900 | 9 | 256 | 212/256/176 | 1.0x1.0x1.2 | 2 |
|  | Rome | 2300/2.86/900 | 9 | 256 | 212/256/176 | 1.0x1.0x1.2 | 2 |
| **Philips** | Groningen | 6.69/3.11/900 | 8 | 270 | 256/232/170 | 1.0x1.0x1.0 | 1.8 |
|  | Zurich | 6.69/3.11/900 | 9 | 270 | 256/232/170 | 1.0x1.0x1.0 | 1.8 |
| **GE** | London | 7.31/3.02/400 | 11 | 270 | 256/256/196 | 1.0x1.0x1.2 | 1.75 |

\* As provided by the manufacturer.

**Figure 1.**
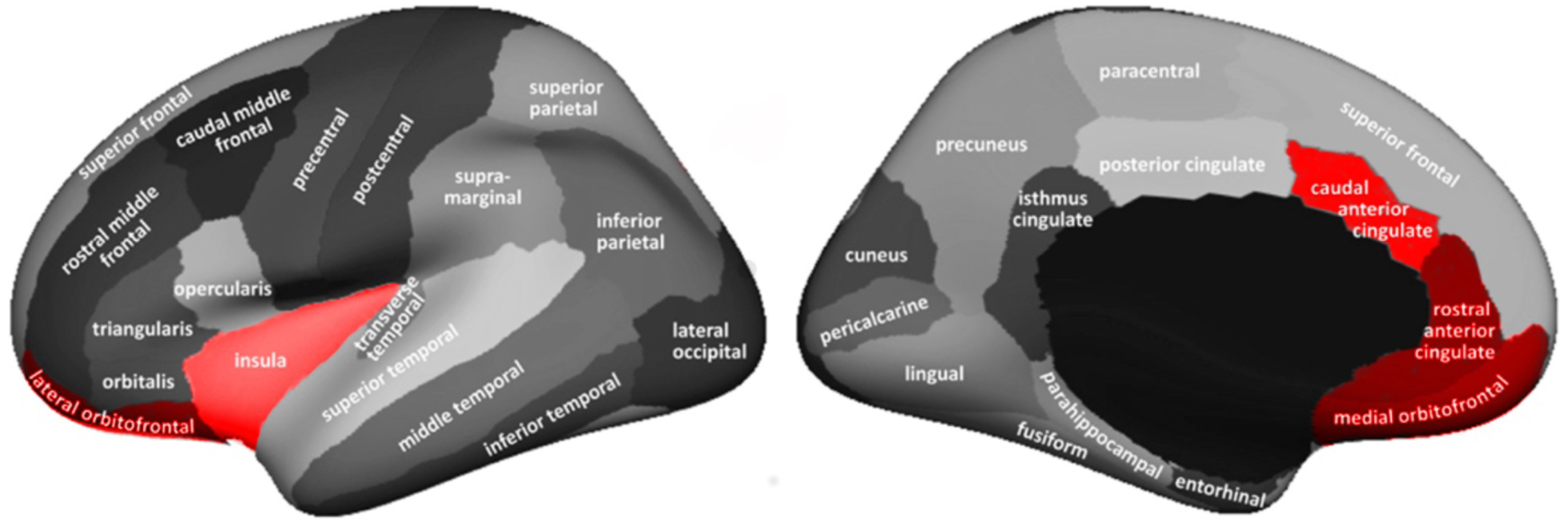
Regions included in the cortical volumes analyses. ACC consisted of rostral and caudal anterior cingulate cortex; vmPFC consisted of lateral-and medial orbitofrontal cortex.

### Quality control

Structural data were all visually inspected and evaluated by an experienced rater (JN). Segmentation of the structures was also visually examined for processing errors and segmentation-accuracy. Since images of participants with externalizing disorders are more prone to motion artefacts, we used a T1 rating system that has been described elsewhere and thoroughly applied to an MRI data-set of children with ADHD and/or CD before.^30^ Scans of 18 participants with ODD/CD and 5 healthy controls were excluded due to anxiety in the scanner or due to poor data quality, which was based on ratings of image sharpness, ringing, contrast to noise ratio of the subcortical structures and of grey and white matter. Total grey matter (GM) volume, total brain volume (TBV), amygdala and ventral striatum volumes are compared across in-and excluded participants in **Figure S1**.

### Statistical analyses

Statistical analyses were performed with the R statistical program.^31^ Group distributions in sex were tested with Pearson’s chi-squared test. Group differences in continuous demographic measures were assessed with one-way analyses of variance (ANOVAs) if assumptions of homogeneity of variance and normality of distributions were met (*p*>0.05 in Levene’s test of homogeneity of variance and Shapiro-Wilk normality test). If these assumptions were violated a non-parametric Kruskal-Wallis rank sum test was used instead. We investigated whether volume of the ventral striatum, amygdala, ACC, vmPFC and insula were related to continuous measures of aggression (proactive and reactive aggression) and to CU-traits and ADHD symptoms by using the multi-level linear mixed effects approach in R with a maximum likelihood fit (nlme package)^32^ with effect sizes being presented as “*r*”. Hemisphere was used as within-subjects’ factor and the respective continuous measure as variable of interest. These analyses were performed in cases only, because of the near-zero scores in controls. Subsequently, sensitivity analyses were performed adding scan-site, age, sex, TBV, IQ, anxiety and total ADHD symptoms to the models (see supplemental material for model statements). Statistical shape analyses based on the vertex-wise F-statistic of the bilateral ventral striatum and amygdala structures were performed using FSL randomise^33^ with 5000 random permutations and threshold-free cluster enhancement (TFCE).^34^ Bonferroni corrections were used for multiple comparisons corrections for testing the shape of multiple structures (*p*_corrected_=0.01). For illustrative purposes, we also performed classical vertex analysis containing vectors displaying the direction of shape alterations.

As part of the supplemental material, we also investigated traditional case-control differences in volume and shape of the same regions as a replication of previous studies^13^ in a larger sample.

## Results

### Demographics

Due to the exclusion of 23 participants, our total sample consisted of 254 participants (n=158 cases and n=96 controls). Out of the 158 cases, 59 were diagnosed with ODD, 11 with CD and 42 with both ODD and CD. The other 46 participants were included as “case” based on a CBCL aggression and/or rule-breaking behavior subscale T-score of ≥ 70. Among the cases, 44 participants were diagnosed with comorbid ADHD. **Table 2** provides a summary of the demographic and clinical information. **Table S1** provides demographic information across the different sites.

**Table 2.**
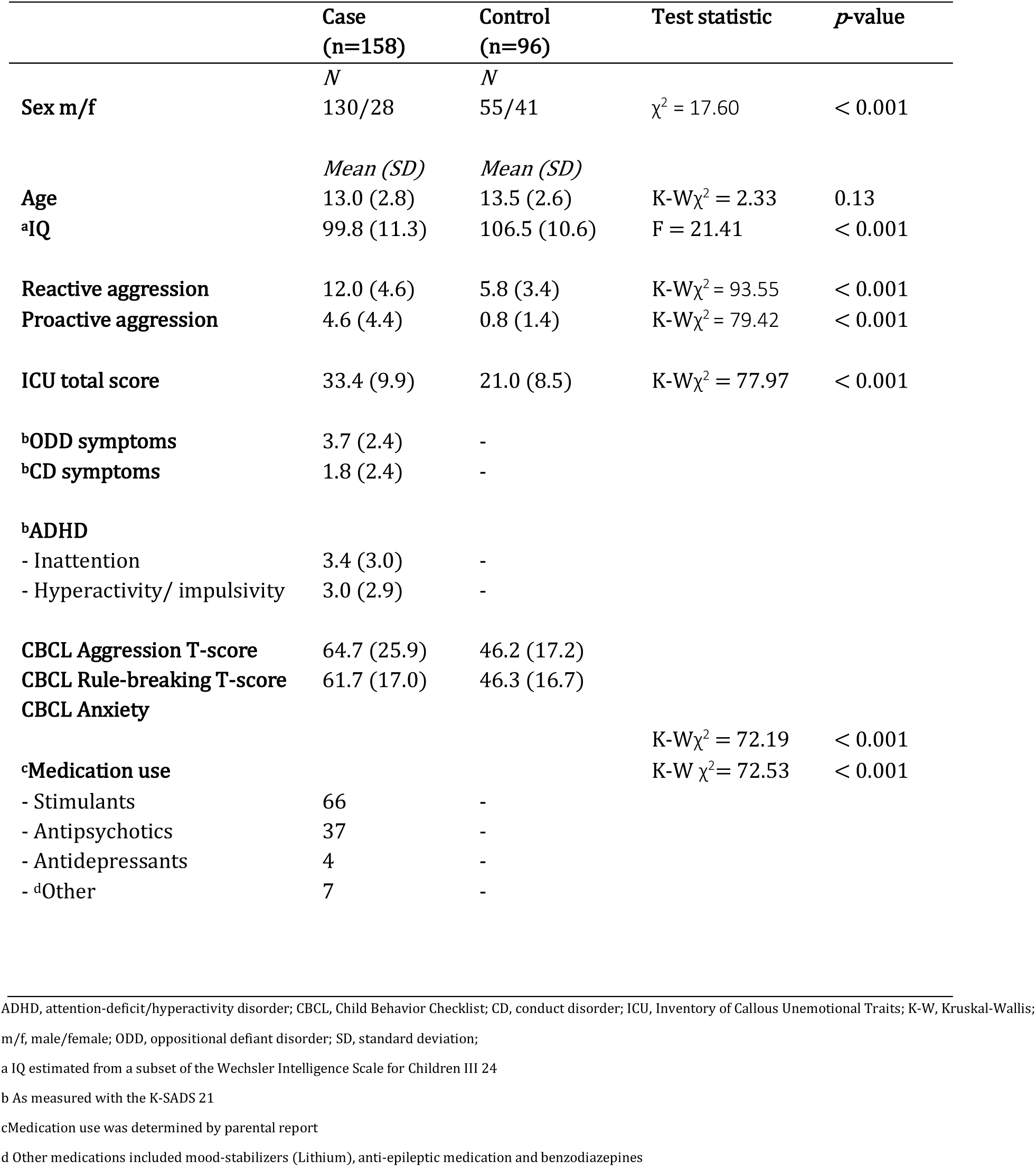
Demographic and clinical characteristics (n=254).

### Continuous measures

In a cases only analysis (n=158), we found an effect of proactive aggression on amygdala volume, showing model improvement from baseline (intercept only) to addition of the continuous variable of proactive aggression (χ^2^(1)=4.17, *p*=0.04). More proactive aggression was associated with smaller amygdala volume (*b*=-11.82, **t**(155)=−1.97, *p*=0.05, *r*=0.15). Reactive aggression was associated with the volume of the insula (χ^2^(1)=5.70, *p*=0.02; *b*=-46.41, **t**(159)=−2.62, *p*=0.01 *r*=0.20), where more reactive aggression were associated with smaller volumes. (see **Figure 2**). These effects were also present when investigating the whole group (n=254; amygdala: *b*=-12.43, **t**(251)=−2.44, *p*=0.02; insula: *b*=-31.90, **t**(251)=−2.65, *p*=0.009), but not in controls only (n=96). No effect of hemisphere or any interaction effects were observed. There was no effect of CU-traits or ADHD symptoms on any of the cortical or subcortical volumes (all *p*-values >0.05). There was also no effect of any of the continuous measures of aggression on ventral striatum or amygdala shape. Group differences are described in the supplemental material.

**Figure 2.**
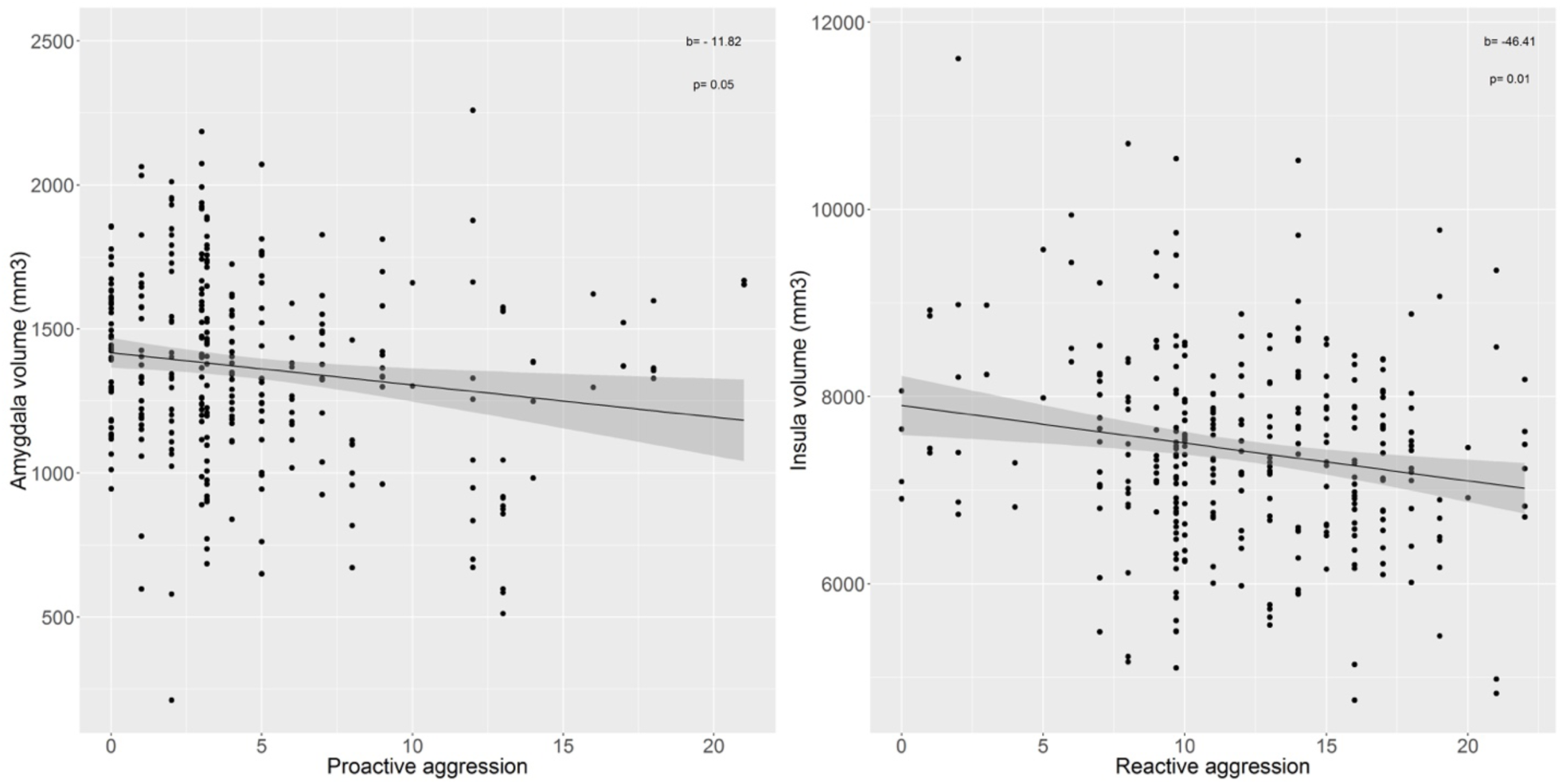
Negative association between proactive aggression and total amygdala volume (A) and between reactive aggression and total insula volume (B) in the case-only analysis.

## Discussion

The current study investigated whether structural brain alterations in regions of interest chosen on the basis of two meta-analyses are differentially associated with reactive and proactive subtypes of aggression. Our main finding is that we could confirm these differential associations by observing reactive aggression to be associated with smaller insula volume and proactive aggression with smaller amygdala volume, although not with the other regions of interest and with relatively small effect sizes. No effects of CU traits or ADHD symptoms were found in any of the regions. In addition, children and adolescents with disruptive behavior showed smaller volumes in the ventral striatum, vmPFC and ACC but not in the amygdala and insula, compared with healthy controls.

Aggression and conduct problems have been associated with several neurocognitive dysfunctions moderated by the presence or absence of psychopathic (CU) traits.^35^ The impaired empathy that underlies psychopathic traits has been linked to more severe forms of aggression (as expressed by high CU traits and/or proactive aggression)^36^, to reduced amygdala response to fear, sadness and pain^37,38^, and to reduced amygdala volume.^11,39,40^ In contrast, heightened threat sensitivity has been associated with reactive aggression and an increased amygdala and insula response to threats and frustration.^41^

Our study confirmed smaller amygdala volumes with an increase in proactive aggression in cases. Despite previous evidence for proactive aggression involving the striatum and vmPFC^8^, related to impaired prediction error signaling and impaired decision making, we did not find an association between proactive aggression and the volumes of these regions in our sample.

Decision making and empathy are highly dependent on each other, where learning and representing the (social) valence of objects and actions is critical in deciding whether to respond empathically, and is related to dysfunction of not only striatum, amygdala and vmPFC, but also insula.^42,43^ In the current study, we found a negative relation between insula volume and reactive aggression, which is associated with responses to threat and frustration. This is in line with studies that report the amygdala to be associated with fear and sadness while the insula is more responsive to angry faces.^44,45^ A reduced insula volume in association with more reactive aggression may explain a deficit in responding to threatening stimuli. Deficits in the insula have also been associated with poor decision making by relating outcome information (reward/punishment) wrongly to responding which in turn increases conduct problems.^7,46,47^

Although no associations with re-or proactive aggression were found, we did report a smaller volume of the ventral striatum, ACC and vmPFC in cases versus controls, supporting a possible deficit related to decision-making in individuals with conduct problems. The reduced volume of the ventral striatum was further supported by a difference in shape, which has not been reported in aggression before and localizes the finding to the anterior part of the ventral striatum. Our findings of reduced vmPFC volume in ODD/CD are in accordance with the idea of reduced empathy, as this region is associated with responses to distress cues.^48^ The decreased volume of the ACC in cases compared with controls further supports the outcome of a meta-analysis.^13^ Atypical responses have been reported in this region in people with conduct problems during negative picture processing.^49^ The ACC has further been implicated in reactive aggression as a defensive response to threat^50^, albeit not further substantiated in our study.

In contrast to previous studies that reported smaller volumes in amygdala and insula in aggressive participants compared to controls, we found associations with continuous measures of proactive and reactive aggression for these regions and no group differences. In addition, we also did not find any association between the volume of any of our regions of interest and the severity of CU traits. Possibly, our group of children and adolescents with ODD and CD showed more reactive (impulsive) forms of aggression and less proactive aggression and CU traits and thus another profile of aggression/conduct problems in comparison to other studies. There was no effect of ADHD symptoms on our main group differences and also not on any of the volumes itself. This is consistent with previous studies.^13,14^

Important strengths of the current study are the investigation of reactive and proactive aggression in a large clinical sample relation to structural brain alterations and the multi-site design. There were also some limitations. First, we included participants with aggression scores in the clinical range on the CBCL who however did not fulfil all diagnostic criteria for ODD and/or CD. Especially the CD group was relatively small. This may have caused heterogeneity in our cases, thus reducing symptom severity and may be reflected in our more profound findings of reduced volume in areas that are less specific for CU traits and lack of empathy types of aggression (ACC, vmPFC and striatum). However, reducing the sample to only cases with a clinical diagnosis did not change our results. Second, the male/female ratio was different for cases and controls which is inherent to the ratio of males/females that are diagnosed with these types of disorders. There was, however, no effect of sex on any of our results. The variance induced by the multi-site setup may have diluted some of our findings. However, investigating in multiple (MRI) centers also allowed us to have a larger sample size compared to many previous studies and facilitated in generalizing our findings.

In conclusion, the current study showed a negative relation between proactive aggression and amygdala volume and between reactive aggression and insula volume and decreased volumes in the ventral striatum, vmPFC and ACC of children and adolescents with ODD/CD compared with controls and. Our findings support the idea of subtype-specific impairments in aggression, where different brain regions are involved in empathy, threat response and decision making which are in turn more associated with either proactive or reactive aggression. This may have implications for designing targeted intervention strategies, which needs to be further explored in future studies.

## Supporting information

## Acknowledgments

This project has received funding from the European Union’s Seventh Framework Programme for research, technological development and demonstration under grant agreement no 602805 (Aggressotype) and 603016 (MATRICS). This work reflects only the authors’ views and the European union is not liable for any use that may be made of the information contained herein. We gratefully acknowledge and thank all the participants and their families for their enthusiastic participation in the study. The authors would also like to thank all other PhD students, post-docs and research assistants for their involvement in data-collection.

T Banaschewski served in an advisory or consultancy role for Actelion, Hexal Pharma, Lilly, Medice, Novartis, Oxford outcomes, PCM scientific, Shire and Viforpharma. He received conference support or speaker’s fee by Medice, Novartis and Shire. He is/has been involved in clinical trials conducted by Shire & Viforpharma. The present work is unrelated to the grants and relationships noted earlier. C Arango has been a consultant to or has received honoraria or grants from Acadia, Ambrosseti, Caja Navarra, CIBERSAM, Fundación Alicia Koplowitz, Forum, Instituto de Salud Carlos III, Gedeon Richter, Janssen Cilag, Lundbeck, Merck, Ministerio de Ciencia e Innovación, Ministerio de Sanidad, Ministerio de Economía y Competitividad, Mutua Madrileña, Otsuka, Roche, Servier, Shire, Schering Plough, Sumitomo Dainippon Pharma, Sunovio and Takeda. D Brandeis serves as an unpaid scientific advisor for an EU-funded Neurofeedback trial unrelated to the present work. JC Glennon has acted as a consultant for Boehringer Ingelheim GmbH. B Franke received an educational speaking fee from Shire and Medice. JK Buitelaar has been consultant to/member of advisory board of and/or speaker for Janssen Cilag BV, Eli Lilly, Bristol-Myer Squibb, Shering Plough, UCB, Shire, Novartis and Servier. He is not an employee of any of these companies, nor a stock shareholder of any of these companies. He has no other financial or material support, including expert testimony, patents, and royalties. The other authors do not report any biomedical financial interests or potential conflicts of interest.

