## Supplementary material for "Reactive/proactive aggression specific cortical and subcortical alterations in children and adolescents with disruptive behavior"

Naaijen et al.

### Details on sites where participants were recruited

Radboud University Medical Center and the Donders institute for Brain, Cognition and Behavior, Nijmegen, The Netherlands; Department of Neuroscience, University Medical Center Groningen, The Netherlands; Central Institute of Mental Health (CIMH), Mannheim, Germany; Department of psychiatry III and Child and Adolescent Psychiatry/Psychotherapy, University of Ulm, Ulm, Germany; Centre for Neuroimaging Sciences, Institute of Psychiatry, Psychology and Neuroscience, King's College London, London, United Kingdom; Department of Child Psychiatry, Institute of Psychiatry, Psychology and Neuroscience, King's College London, London, United Kingdom; Department of Child and Adolescent Psychiatry and Psychology, Neurosciences Institute, Hospital Clinic de Barcelona, Barcelona, Spain; Hospital Gregorio Marañón, Madrid, Spain; MR Center, Psychiatric University Hospital, Zurich, Switzerland; IRCCS Santa Lucia Foundation, Rome, Italy

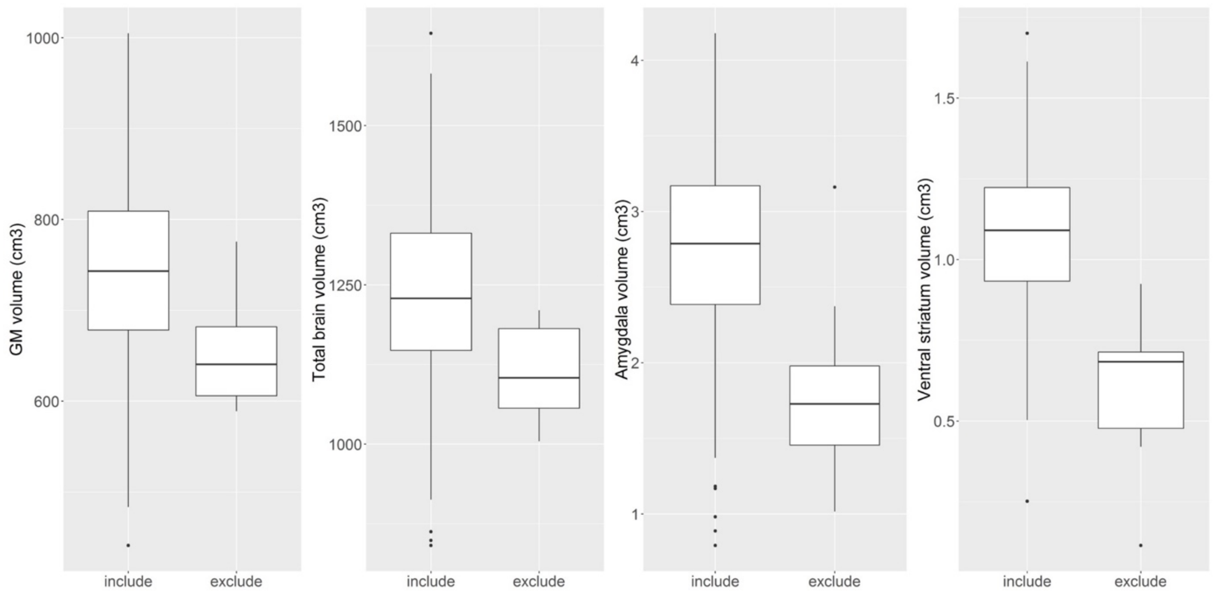


**Figure S1.** Differences in GM volume, total brain volume, amygdala and ventral striatum volume for participants that were included versus participants that were excluded based on the rating system of

Backhausen et al. 2016. All volume differences were significant ($p$-values $<0.05$) between in- and excluded participants and no inclusion/exclusion by diagnosis interactions were found (all $p$-values $> 0.05$).

**Details of statistical analyses and model statement**

For each analysis considering associations with continuous measures of aggression and group differences in volume (or thickness/surface area) we used the following R codes using a multi-level linear mixed effects approach considering dependency in the data with a 2 level-hierarchy of nesting of the within subject's variable *hemisphere* in *subject* (using the nlme package; Pinheiro et al. 2007):

| | |
|---|---|
| Baseline: | Volume ~ 1, random = ~1\|subject/hemisphere, method="ML" |
| Hemisphere: | Volume ~ 1 + hemisphere, random = ~1\|subject/hemisphere, method="ML" |
| Diagnosis: | Volume ~ 1+ hemisphere + diagnosis**, random = ~1\|subject/hemisphere, method="ML" |
| H×D: | Volume ~ 1+ hemisphere + diagnosis + hemisphere:diagnosis, random = ~1\|subject/hemisphere, method="ML" |

** the variable "diagnosis" can be replaced with continuous measures of reactive aggression, proactive aggression, CU traits and ADHD symptoms using the same model

Anova (Baseline, Hemisphere, Diagnosis, H×D)[a]

Summary(H×D)[b]

[a] Model improvement from baseline to addition of diagnosis and addition of hemisphere (and their possible interaction) was described by the likelihood ratio ($\chi^2$).

[b] If the likelihood ratio was significant, the linear mixed effects model itself was interpreted. For the continuous measures of proactive aggression, reactive aggression and cu-traits we used similar models. The effect of confounders was considered separately by subsequently adding age, sex, scan-site, TBV, IQ and ADHD-symptoms to the above described models.

### Analyses of traditional case-control differences

In addition to associations between (sub)cortical volumes and continuous measures of aggression, we investigated traditional case-control differences to replicate previous studies' results[3] in a larger sample. Since our group of "cases" consisted not only of participants with a clinical diagnosis, we performed a sensitivity analysis for the group differences in those with an ODD/CD diagnosis (n=112 cases vs n=96 controls). Additionally, a sensitivity analysis was performed for the subgroup that did not use any medication (n=61 cases vs n=96 controls). Sensitivity analyses are described in detail in the supplemental information.

The effect of ADHD was both included in the sensitivity analyses to account for the effect of ADHD in possible group differences and to explore the effect of ADHD symptoms as a factor in itself. When volumetric differences were found, CT and SA were considered separately for the cortical regions. Corrections for multiple comparisons in the categorical group-analyses were based on the effective number of independent tests[4], resulting in an adjusted alpha level of 0.01 (based on 3.8 effective tests). Statistical shape analyses based on the vertex-wise F-statistic of the bilateral ventral striatum and amygdala structures were performed using FSL randomise[5] with 5000 random permutations and threshold-free cluster enhancement (TFCE).[6] Bonferroni corrections were used for multiple comparisons corrections for testing the shape of multiple structures ($p_{corrected}$=0.01). For illustrative purposes, we also performed classical vertex analysis containing vectors displaying the direction of group differences.

### Results of traditional case-control differences

Cases and controls differed significantly in bilateral ventral striatum, ACC and vmPFC volumes, showing model improvement from baseline (intercept only) to addition of the between-subject variable diagnosis (ventral striatum: $\chi^2(1)=5.93$, $p=0.01$; ACC: $\chi^2(1)=4.13$, $p=0.04$; vmPFC: $\chi^2(1)=9.05$, $p=0.003$). In all regions, cases had significantly smaller total volumes than controls ($b$=-43.72, **t**(251)=-2.98, $p$=0.003, $r$=0.18; $b$ =-358.52, **t**(251)=-2.54, $p$=0.01, $r$=0.16; $b$=-772.31, **t**(251)=-2.96, $p$=0.003, $r$=0.18 for ventral striatum, ACC and vmPFC respectively).

In the analysis of the smaller sub-group of cases with a clinical diagnosis of ODD and/or CD (n=112) or the sub-group not using any medication (n=61), group differences in ventral striatum, ACC and vmPFC remained to be present (all $p$-values <0.05). The exception was that the case-control difference in the ACC lost statistical significance when including only unmedicated cases.

Considering the CT and SA of the ACC and vmPFC separately did not reveal any significant group differences ($p$-values > 0.05). Shape analyses of the subcortical regions revealed differences in the left (but not right) ventral striatum (**Figure S2**). These results were in accordance with the reductions found in volume, showing an overall inward position of the vertices (corrected *p*-value of 0.005) with the largest difference in one anterior cluster of voxels (voxel-coordinates: x=102, y=143, z=69, size=28 voxels). This showed that the smaller volume in cases compared with controls was specific for the anterior part of the ventral striatum. No shape differences between ODD/CD and controls were found for the amygdala.

**Sensitivity analyses**

When checking the observed associations for the effects of age, sex, TBV, IQ, anxiety, total ADHD symptoms and scan-site, the association between proactive aggression and amygdala volume remained significant for all possible confounders (all *p*-values $< 0.05$) except for scan-site ($b$=-2.79, **t**(147)=-0.58, $p$=0.56). The association between reactive aggression and insula volume survived all sensitivity analyses (all *p*-values remained $< 0.05$). When repeating the analyses in male cases only (n=128), the association between proactive aggression and amygdala volume and reactive aggression and insula volume remained significant (both *p*-values $< 0.05$).

The between-group differences remained significant in the ventral striatum for all of the possible confounders (all *p*-values remained $<0.05$). For the ACC, the effect of diagnostic status remained significant for all confounders except for scan-site ($b$=-201.20, **t**(243)= -1.49, $p$ =0.14) and anxiety ($b$=-201.01, **t**(227)= -1.23, $p$=0.22). For the vmPFC, adding total ADHD symptoms to the model caused the effect of diagnosis to become trend-level ($b$=-539.77, **t**(250)=-1.71, $p$=0.09). Age was not of influence on ventral striatum or ACC volume, but negatively affected the vmPFC volume, where older age was associated with lower volume ($\chi^2(1)$=4.93, $p$=0.03; $b$=-100.49, **t**(250) = -2.22, $p$=0.03, $r$=0.14). In all three regions, males showed higher volumes than females (ventral striatum: $\chi^2(1)$=6.03, $p$=0.01; $b$=37.12, **t**(250) = 2.46, $p$=0.01, $r$=0.15; ACC: $\chi^2(1)$=12.70, $p<0.001$; $b$=480.06, **t**(250) = 3.59, $p<0.001$, $r$=0.22; vmPFC: $\chi^2(1)$=42.72, $p<0.0001$; $b$= 1789.91, **t**(250) = 6.79, $p<0.0001$, $r$=0.39). TBV was positively associated with regional brain volume in ventral striatum ($\chi^2(1)$=74.70, $p<0.0001$; $b$=0.00039, **t**(250) = 9.28, $p<0.0001$, $r$=0.51), ACC ($\chi^2(1)$=137.12, $p<0.0001$; $b$=0.0045, **t**(250) = 13.42, $p<0.0001$, $r$=0.65), as well as vmPFC ($\chi^2(1)$=194.40, $p<0.0001$; $b$=0.016, **t**(250) = 17.02, $p<0.0001$, $r$=0.73). No effect of IQ, anxiety or ADHD symptoms was found on any of the regions. The effect of scan-site on the ventral striatum, ACC and vmPFC volume is shown and described in **Figure S3**.

**Table S1.** Demographic characteristics across sites

| | | Case (n=158) | Control (n=96) | Ratio cases (dark grey) /controls (light grey) |
|---|---|---|---|---|
| **Nijmegen (n=56)** | | | | |
| | Age | 13.5 (2.9) | 13.3 (2.6) | 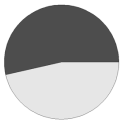 |
| | IQ | 99.0 (12.6) | 107.5 (13.0) | |
| | Sex m/f | 20/10 | 18/8 | |
| **Groningen (n=21)** | | | | |
| | Age | 14.3 (3.0) | 12.2 (2.4) | 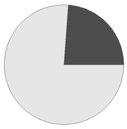 |
| | IQ | 99.9 (11.3) | 102.2 (1.9) | |
| | Sex m/f | 13/3 | 1/4 | |
| **Mannheim (n=32)** | | | | |
| | Age | 12.8 (2.4) | 13.1 (3.2) | 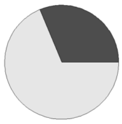 |
| | IQ | 102.2 (10.1) | 115.1 (7.0) | |
| | Sex m/f | 19/3 | 8/2 | |
| **Ulm (n=20)** | | | | |
| | Age | 10.4 (1.9) | 13.6 (3.3) | 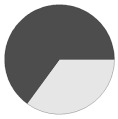 |
| | IQ | 100.7 (16.5) | 101.4 (8.0) | |
| | Sex m/f | 7/0 | 4/9 | |
| **London (n=29)** | | | | |
| | Age | 13.4 (2.9) | 13.7 (2.1) | 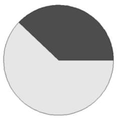 |
| | IQ | 94.9 (10.6) | 111.8 (11.0) | |
| | Sex m/f | 18/0 | 8/3 | |
| **Barcelona (n=34)** | | | | |
| | Age | 12.4 (2.9) | 14.7 (2.3) | 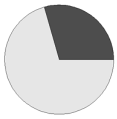 |
| | IQ | 101.7 (11.1) | 105.4 (6.3) | |
| | Sex m/f | 18/6 | 5/5 | |
| **Madrid (n=22)** | | | | |
| | Age | 14.2 (2.1) | 14.7 (1.8) | 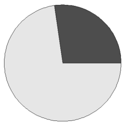 |
| | IQ | 98.0 (9.1) | 105.7 (9.3) | |
| | Sex m/f | 12/3 | 5/2 | |
| **Zurich (n=25)** | | | | |
| | Age | 11.0 (2.0) | 11.7 (1.5) | 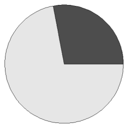 |
| | IQ | 102.9 (11.6) | 99.6 (7.9) | |
| | Sex m/f | 14/4 | 3/4 | |
| **Rome (n=15)** | | | | |
| | Age | 14.0 (2.4) | 16.1 (1.8) | 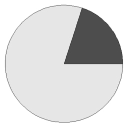 |
| | IQ | 94.1 (9.7) | 97.2 (2.8) | |
| | Sex m/f | 11/1 | 1/2 | |

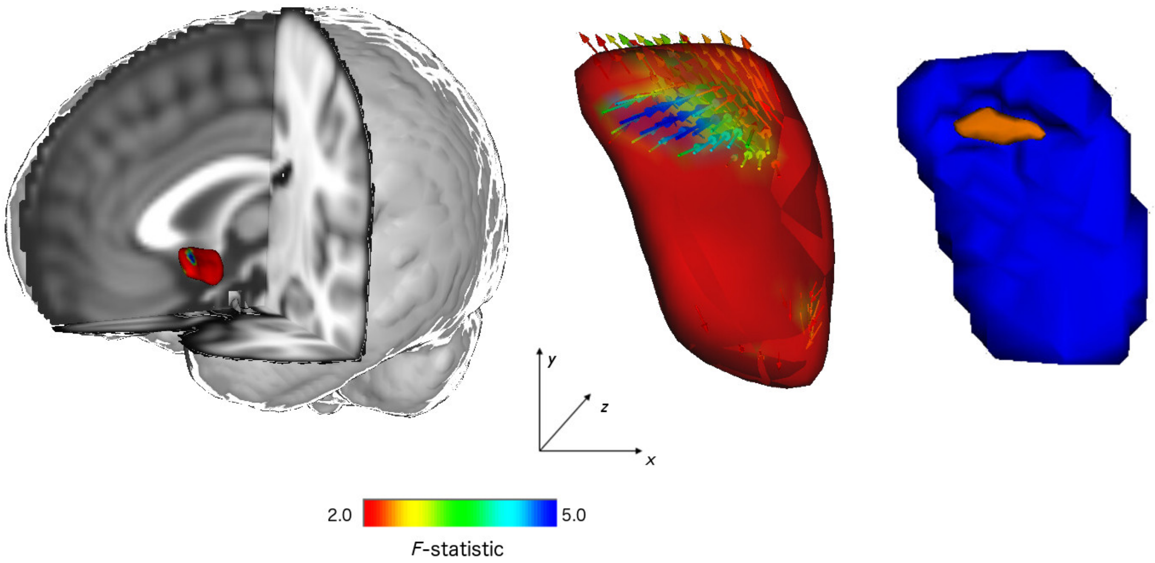


**Figure S2.** Vertex analyses of shape alterations in the left ventral striatum. Left panel shows the anatomical location of the area and the local areas exhibiting shape change (alterations in color). The middle panel shows shape changes in the ODD/CD participants compared with controls with vector directions. Inward direction represents relative inward position of the vertices, pointing to regional decreased shapes. The color of the surface and the arrows indicate the Pillai's trace F-statistic. The right panel shows the results analyzed with the vertex-wise F-statistic. The region in orange corresponds to the anterior part of the ventral striatum shown to be smaller in ODD/CD than in controls (which resembles the dark blue arrows in the middle panel).

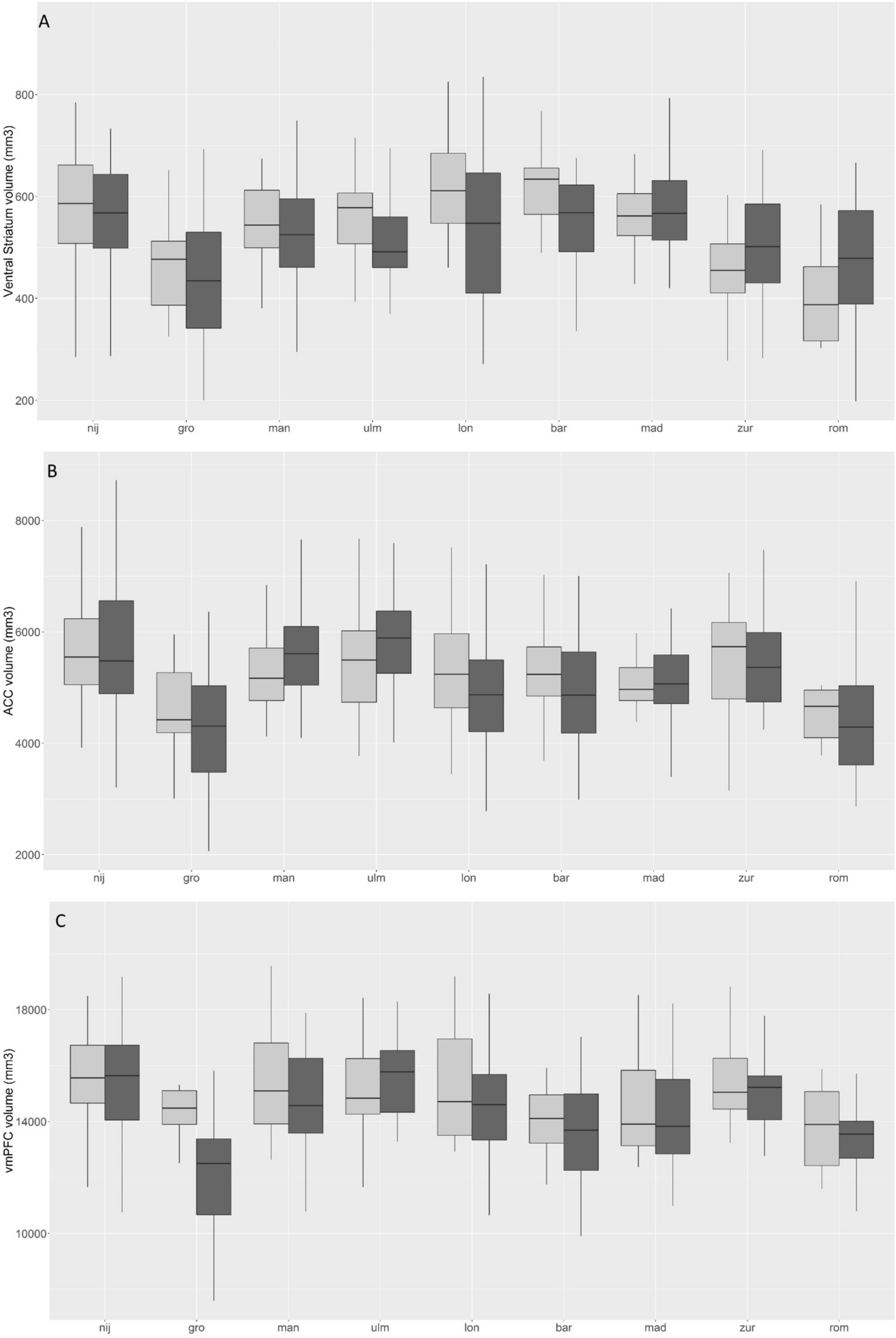

A
Ventral Striatum volume (mm3)
800
600
400
200
nij
gro
man
ulm
lon
bar
mad
zur
rom
B
ACC volume (mm3)
8000
6000
4000
2000
nij
gro
man
ulm
lon
bar
mad
zur
rom
C
vmPFC volume (mm3)
18000
14000
10000
nij
gro
man
ulm
lon
bar
mad
zur
rom

**Figure S2.** Effect of scan-site on ventral striatum volume (A), ACC volume (B) and vmPFC volume (C) split by diagnosis (light-grey = controls, dark-grey = cases). Note: figure displays raw values and SEs. Site was influencing ventral striatum volume ($\chi^2(8)=47.68$, $p<0.0001$). Breaking this down in contrasts showed Groningen, Zurich and Rome to have lower volumes than the reference-site Nijmegen (Groningen: $b=-124.54$, $\mathbf{t}(243)=-5.03$, $p<0.0001$, $r=0.31$; Zurich: $b=-68.16$, $\mathbf{t}(243)=-2.94$, $p=0.004$, $r=0.19$; Rome: $b=-99.76$, $\mathbf{t}(243)=-3.46$, $p<0.001$, $r=0.22$). Scan-site was additionally influencing ACC volume ($\chi^2(8)=53.42$, $p<0.0001$), where Groningen ($b=-1294.13$, $\mathbf{t}(243)=-5.90$, $p<0.0001$, $r=0.36$), London ($b=-587.79$, $\mathbf{t}(243)=-3.02$, $p=0.003$, $r=0.19$), Barcelona ($b=-612.83$, $\mathbf{t}(243)=-3.29$, $p=0.001$, $r=0.21$), Madrid ($b=-476.27$, $\mathbf{t}(243)=-2.22$, $p=0.03$, $r=0.14$) and Rome ($b=-1170.11$, $\mathbf{t}(243)=-4.58$, $p<0.0001$, $r=0.28$) showed smaller volumes than the reference site Nijmegen. Scan-site influenced vmPFC volume as well ($\chi^2(8)=47.29$, $p<0.0001$). Groningen ($b=-2723.77$, $\mathbf{t}(243)=-5.86$, $p<0.0001$, $r=0.35$), Barcelona ($b=1630.36$, $\mathbf{t}(243)=-4.13$, $p<0.0001$, $r=0.26$), Madrid ($b=-1119.76$, $\mathbf{t}(243)=-2.46$, $p=0.01$, $r=0.16$) and Rome ($b=-1909.16$, $\mathbf{t}(243)=-3.52$, $p<0.001$, $r=0.22$) showed smaller volumes than the reference site Nijmegen. Removal of the most influential site Groningen did not influence our main effects of interest of diagnosis on volume ($p=0.007$, $p=0.04$ and $p=0.03$ for ventral striatum, ACC and vmPFC respectively). Nij, Nijmegen; gro, Groningen; man, Mannheim; ulm, Ulm; lon, London; bar, Barcelona; mad, Madrid, zur, Zurich, rom, Rome.
